# Secreted filarial nematode galectins modulate host immune cells

**DOI:** 10.1101/2022.05.23.493127

**Authors:** Hannah J. Loghry, Noelle A Sondjaja, Sarah J Minkler, Michael J Kimber

## Abstract

Lymphatic filariasis (LF) is a mosquito-borne disease caused by filarial nematodes including *Brugia malayi*. Over 860 million people worldwide are infected or at risk of infection in 72 endemic countries. The absence of a protective vaccine means that current control strategies rely on mass drug administration programs that utilize inadequate drugs that cannot effectively kill adult parasites, thus established infections are incurable. Progress to address deficiencies in the approach to LF control is hindered by a poor mechanistic understanding of host-parasite interactions, including mechanisms of host immunomodulation by the parasite, a critical adaptation for establishing and maintaining infections. The canonical type 2 host response to helminth infection characterized by anti-inflammatory and regulatory immune phenotypes is modified by filarial nematodes during chronic LF. Current efforts at identifying parasite-derived factors driving this modification focus on parasite excretory-secretory products (ESP), including extracellular vesicles (EVs). We have previously profiled the cargo of *B. malayi* EVs and identified *B. malayi* galectin-1 and galectin-2 as among the most abundant EV proteins. In this study we further investigated the function of these proteins. Sequence analysis of the parasite galectins revealed highest homology to mammalian galectin-9 and functional characterization identified similar substrate affinities consistent with this designation. Immunological assays showed that Bma-LEC-2 is a bioactive protein that can polarize macrophages to an alternatively activated phenotype and selectively induce apoptosis in Th1 cells. Our data shows that an abundantly secreted parasite galectin is immunomodulatory and induces phenotypes consistent with the modified type 2 response characteristic of chronic LF infection.

## 1. Introduction

Lymphatic filariasis (LF) is a mosquito-borne neglected tropical disease (NTD) caused by parasitic, filarial nematodes including *Brugia malayi, Brugia timori,* and *Wuchereria bancrofti.* LF is endemic in 72 countries and over 860 million people are infected or at risk of infection (1). Infection is often asymptomatic, but can also result in clinical symptoms with extreme morbidity leading to mortality in some cases. Morbidities of this disease include lymphangitis, lymphedema, primarily in the extremities, and secondary bacterial infection/dermatitis (2). Mass drug administration programs are the most common disease control strategies employed in endemic countries. With the absence of a protective vaccine, these programs rely on inadequate drugs that cannot effectively kill adult parasites leaving established infections incurable. While solutions to both these deficiencies are being investigated, progress is hindered by a poor mechanistic understanding of parasite biology and host-parasite interactions including mechanisms of host immunomodulation by the parasite.

Host immunomodulation is critical for establishing and maintaining infections. In chronic LF, this modulation is seen in the development of a “modified” type 2 immune response that is characterized by an increase in alternatively activated macrophage (3–5) and regulatory T cell populations (5–10), an increase in the IL-4 and IL-10 regulatory environment (11–13), and a suppression, or hyporesponsiveness, of effector T cells (14–17). The outcome of this modification is to create a state of immune tolerance where the host can maintain an active immune response, but damage to the parasite is limited. This modified type 2 response exists, at least in part, as an immune evasion strategy with parasite excretory-secretory products (ESP) as a well-established source of effector molecules driving the modifications. Parasite ESP encompass freely secreted proteins and nucleic acids, as well as extracellular vesicles (EVs), a heterogenous group of cell-derived, membrane-bound vesicles that are known to be involved in various physiological processes and inter-cell communication (18, 19). It has been identified that filarial nematodes secrete EVs and that they contain cargo such as proteins, lipids and small RNA species some with immunomodulatory functions (20–25). In addition, there is evidence that parasitic nematode EVs are involved in the immunomodulation of the mammalian host (20,21,25–32). EVs represent a compelling mechanism for the non-canonical secretion of immunomodulatory molecules and their subsequent protected trafficking and delivery to host cells but we have a poor understanding of how cargo molecules contained within EVs might exert their modulatory effects.

Our lab has previously identified two parasite derived proteins, Bma-LEC-1 and Bma-LEC-2, as amongst the most abundant proteins found within EVs secreted by adult female *B. malayi*. Galectins are a protein family defined by their conserved β-galactoside binding sites within the approximate 130 aa carbohydrate recognition domain (CRD) (33). This family is divided into three sub-domains according to their CRD sequence and structure. The prototypical galectins contain one CRD domain, the chimeric galectin contain a single CRD domain connected to a non-lectin N-terminal region, and the tandem-repeat type galectins contain two CRD domains separated by a linker peptide (34). Synthesis of galectins occurs in the cytosol from where they can be directed to various cell compartments including vesicular trafficking pathways but not the ER-Golgi secretory pathway; as galectins do not contain a N-terminal signal sequence they must be secreted through alternative mechanisms such as EVs (35). The functional effects of galectins originate from their galactose-containing glycan binding properties and their ability to oligomerize. Tandem-repeat type galectins can oligomerize from associations of a N-terminal CRD with a N-terminal CRD of another galectin or a C-terminal CRD with a C-terminal CRD of another galectin (36–38). The multivalency created by these galectin multimers, and by the presence of multiple glycans on a glycoprotein, can cause crosslinking of glycoproteins leading to glycoprotein segregation, formation of lattices, or endocytosis of bound glycoproteins (36,38– 41). In addition, galectins have been shown to bind cytosolic and nuclear, non-carbohydrate ligands but the mechanisms of this are poorly understood (42, 43). It must be noted that there are no true *a priori* functions of galectins, their functions depend on the effect that the specific glycoprotein or glycolipid counterreceptors has on a particular cell in a particular context (44). That said, mammalian galectins are known to have diverse and complex effects on the immune system, modulating both myeloid and lymphoid cells to elicit, in general, immunosuppressive responses (reviewed (44, 45).

We hypothesize that the parasite-derived proteins, Bma-LEC-1 and Bma-LEC-2, found within secreted EVs, are effector molecules that phenocopy host galectins to directly modulate the host immune response. To test this hypothesis, we produced recombinant Bma-LEC-1 and Bma-LEC-2 and investigated their glycan binding properties. We identified that Bma-LEC-1 and Bma-LEC-2 are most closely related to and have similar binding affinities as mammalian galectin-9. A human monocyte cell line and primary murine T cells were used as a model to further investigate the immunosuppressive functions of Bma-LEC-1 and Bma-LEC-2. It was found that although Bma-LEC-1 was not bioactive in the assays used here, treatment of human macrophages with Bma-LEC-2 lead to an increase in expression and production of IL-10, a potent anti-inflammatory cytokine, whilst Bma-LEC-2 also selectively induced apoptosis in Th1 cells, but not naïve CD4^+^ T cells. These findings provide evidence that parasite derived galectins found within extracellular vesicles are capable of modulating the host immune response and promoting the modified type 2 immune phenotypes seen in chronic filarial infection.

## 2. Materials and Methods

### 2.1 *Brugia malayi* Maintenance

*B. malayi* were cultured and maintained as previously described (24), briefly, *B. malayi* parasites were obtained from the NIH/NIAID Filariasis Research Reagent Resource Center (FR3) at the University of Georgia, USA. Persistent *B. malayi* infections at FR3 are maintained in domestic short-haired cats. To obtain adult stage *B. malayi*, jirds were infected intraperitoneally with approximately 400 L3 stage parasites. 120 days post infection jirds were necropsied to collect adult stage parasites. L3 stage *B. malayi* were obtained from dissection of anesthetized *Aedes aegypti* 14 days post-infection. Microfilaria stage *B. malayi* were obtained from a lavage of the peritoneal cavity of a euthanized gerbil. Upon receipt at ISU, all *B. malayi* parasites were washed several times in worm culture media warmed to 37°C (RPMI with 1% HEPES, 1% L-glutamine, 0.2% Penicillin/Streptomycin, and 1% w/v glucose (Thermo Fisher Scientific)), counted, and cultured at 37°C with 5% CO_2_.

### 2.2 Expression and Purification of *B. malayi* Galectins

Total RNA was isolated from adult female *B. malayi*. Briefly, approximately 30 parasites were homogenized in Trizol (Thermo Fisher Scientific) and mixed with chloroform (0.2 ml chloroform per ml Trizol) (Sigma-Aldrich, St. Louis, MO). Samples were shaken vigorously for 20 s and allowed to sit at room temperature for 3 min followed by centrifugation at 10,000 x *g* for 18 min at 4°C. The aqueous phase was collected, and an equal volume of 100% ethanol was added. RNA was then purified and collected using a RNeasy Mini Kit (Qiagen, Hilden, Germany) according to manufacturer’s instructions. cDNA synthesis was performed using the Superscript III first strand cDNA synthesis kit (Thermo Fisher Scientific) per manufacturer’s protocols. Gene sequences for *Bma-LEC-1* (WBGene00226528) and *Bma-LEC-2* (WBGene00224538) were obtained from wormbase parasite (46, 47). Primers were designed for the predicted coding region of Bma-LEC-1 and Bma-LEC-2 that incorporated restriction digest sites facilitating recombination into the pOETIC 6xHis Transfer Plasmid (Mirus Bio, Madison, WI) (Supplemental Materials 1). The complete coding sequence for each galectin was PCR amplified from cDNA using Platinum Taq Polymerase High Fidelity (Thermo Fisher Scientific). The PCR product was visualized through TAE agarose gel electrophoresis and gel purified with Purelink Quick Gel Extraction kit (Thermo Fisher Scientific) per manufacturer’s instructions.

Bma-LEC-1 and -2 PCR products were digested and ligated into pOET1C 6xHis Transfer plasmid using T4 DNA ligase (Thermo Fisher Scientific), transformed into ampicillin resistant NEB alpha competent *E. coli* cells (New England Biolabs, Ipswich, MA), inoculated into LB + Ampicillin media (10g/L Tryptone, 5g/L Yeast Extract, 10g/L NaCl, 100 µg/mL Ampicillin) (Sigma-Aldrich) and incubated at 200 rpm at 37°C overnight. Plasmid was purified from transformed *E.coli* using Genelute Endotoxin-free plasmid midiprep kit (Sigma-Aldrich) per manufacturers’ instructions and sequenced to confirm insert fidelity and orientation. The *Spodoptera frugiperda*-derived Sf21 cell line (Thermo Fisher Scientific, Waltham, MA) was maintained in Insect-XPRESS cell culture media (Lonza Bioscience, Basel, Switzerland) supplemented with 10% Chrysalis insect cell qualified FBS (Gemini Bio Products, West Sacramento, CA), 1% Penicillin (10,000 U/ml) 1% Streptomycin (10,000 ug/ml), and 0.25 µg/ml Amphotericin B (Thermo Fisher Scientific) in normoxic conditions at 28°C. Positive recombinant plasmid was transfected into Sf21 cells using the Flashbac Ultra Baculovirus Expression System (Mirus Bio). Viral Titers were determined with the BacPak qpcr Titration kit (Takara Bio, Kusatsu, Shiga, Japan). Infected Sf21 cells were collected and lysed in dPBS (Thermo Fisher Scientific) containing Halt Protease Inhibitor Cocktail (Thermo Fisher Scientific) using a disruptor genie (Scientific Industries, Bohemia, NY). Protein expression was confirmed by western blot using a 6x His-Tag HRP conjugated Monoclonal Antibody (Thermo Fisher Scientific). Once verified, mass production of recombinant protein was achieved through large scale infections of 400 mL of Sf21s cells at a multiplicity of infection of 8. Cells were incubated for 3 days at 28°C at 130 rpm post infection.

To purify recombinant protein, cell pellets were lysed in dPBS containing protease inhibitors (Thermo Fisher Scientific) using glass beads and a cell disruptor genie (Scientific Industries). Cells were disrupted for 2 min then incubated on ice for 2 min, repeating for a total of 7 rounds of disruption. Disrupted cells were then centrifuged at 15,800 x *g* for 30 min at 4°C and the resulting supernatant collected with an initial protein purification performed using the HisPur Ni-NTA Resin (Thermo Fisher Scientific) following manufacturer’s instructions for native protein confirmation and gravity flow columns. Eluted proteins were further purified on a HiPrep Sephacryl S-200 HR size exclusion column (Cytiva, Marlbourough, MA) on a AKTA Pure FPLC (Cytiva). FPLC fractions were validated for purity using an SDS-PAGE gel stained with Coomassie Blue. Only clean fractions devoid of debris or non-specific proteins and containing the protein of interest were used for downstream experimentation. Double-purified protein was concentrated with Amicon Ultra Centrifugal Filters according to manufacturer’s instructions (Sigma-Aldrich) and the concentration determined by a BCA assay using the Pierce BCA Protein assay kit (Thermo Fisher Scientific). Protein samples were aliquoted and stored at -80°C for future use.

### 2.3 Assessment of glycan binding properties

The functionality of the recombinant *B. malayi* galectins was first analyzed by hemagglutination assay as described by Sano and Ogawa (2014) (48). Briefly, two-fold serial dilutions of recombinant Bma-LEC-1 and Bma-LEC-2 were prepared in V-bottom 96-well plates (Greiner Bio-One, Kremsmünster,Austria) followed by the addition of a 4% solution of trypsinized, glutaraldehyde-fixed, rabbit erythrocytes (Innovative Research, Novi, MI) prepared as previously described (48). Briefly, 5 mL of isolated erythrocytes were suspended in 100 mL of a 0.1% (w/v) trypsin solution and incubated at 37°C for 1 h. After, erythrocytes were centrifuged at 500 x *g* for 5 min and washed in 50 mL of dPBS (Thermo Fischer Scientific). Centrifuged erythrocytes were resuspended in 25 mL of a 1% glutaraldehyde solution in dPBS and incubated for 1 h at room temperature with a gentle shaking. Following, erythrocytes were washed twice with 25 mL of 0.1 M glycine solution in dPBS. Erythrocytes were washed a final time in 25 mL of dPBS and stored as a 10% (v/v) erythrocyte suspension in dPBS at 4°C for future use. Plates were scored for hemagglutination activity after 30 min. As described previously (48), a solid colored well (mat) indicates that the galectin can agglutinate the erythrocytes by binding to the carbohydrate moieties on the surface of the cells and a dot in the center of the well indicates that the galectin is inactive or an inhibitor is present.

This same basic assay was also used to further investigate carbohydrate binding specificity of Bma-LEC-1 and Bma-LEC-2. An initial screening of the ability of recombinant Bma-LEC-1 and Bma-LEC-2, human galectin-9, and mouse galectin-9 to bind to 1 M galactose, 900 mM N-acetylgalactosamine (GalNAc), 1M lactose, 32.6mM N-acetyllactosamine (LacNAc), 2 M mannose, and 2 M glucose was tested. To determine the affinity of these proteins to the various β-galactosides serial dilutions of galactose, N-acetylgalactosamine, lactose, N-acetyllactosamine, mannose, and glucose were prepared, followed by addition of the galectin of interest. The protein carbohydrate mixture was allowed to incubate at room temperature for 30 min after which a 4% rabbit erythrocyte solution was added. The reaction was allowed to proceed for an additional 30 min before scoring. In general, a solid colored well indicates that the galectin present is not binding to the carbohydrate of interest but is instead binding the erythrocytes forming a lattice of erythrocytes on the bottom of the well. A well with a dot in the center indicates that the galectin present is binding the carbohydrate of interest, not the erythrocytes, allowing them to sediment at the bottom of the well.

A more thorough assessment of parasite galectin activity was conducted by glycan binding array, performed by the Protein-Glycan Interaction Resource of the Consortium for Functional Glycomics (CFG) and the National Center for Functional Glycomics (NCFG) at Beth Israel Deaconess Medical Center, Harvard Medical School (supporting grants R24 GM137763). Briefly, samples were assayed against a comprehensive array of 584 glycans provided by the NCFG. The array was generated from a library of natural and synthetic mammalian glycans with amino linkers printed onto N-hydroxysuccinimide (NHS)-activated glass microscope slides forming covalent amide linkages (49). The glycan spotting concentration was 100 μM printed in 6 technical replicates on each microarray. Alexa Fluor 488-labeled rBma-LEC-1 and rBma-LEC-2 were generated using the Alexa Fluor 488 Microscale Protein Labeling Kit per manufacturer’s instructions (Thermo Fischer Scientific) with 5 μg/ml and 50 μg/ml concentrations incubated on the array for 1 h at room temperature. After washing off unbound sample with successive washes of PBS-tween, PBS, and water, slides were dried and scanned on a GenePix Microarray scanner (Molecular Devices, San Jose, CA, USA) at 488 nm. The Glycan Array Dashboard software (50) was used to compare binding specificities between galectins. GlycoGlyph (51), a glycan drawing program, was used to visualize glycan structures.

### 2.4 RT-qPCR analysis of galectin gene expression

30 adult female, 30 adult male, 1,500 L3 and 2x10^6^ microfilariae life stage *B. malayi* were manually homogenized in Trizol (Thermo Fischer Scientific) using a mortar and pestle and total RNA extracted as described above. cDNA was synthesized using a Superscript III First Strand cDNA Synthesis Kit (Thermo Fisher Scientific) according to manufacturer’s instructions. RT-qPCR was conducted using SYBR green (Thermo Fisher Scientific) with gene-specific primers designed against *Bma-lec-1* and *Bma-lec-2* transcripts (Supplemental Materials 1). Average C_T_ values for each life stage cDNA were recorded and relative abundance to the housekeeping gene *B. malayi NADH Dehydrogenase subunit 1* (ND1) (NC_004298.1) using ΔΔC_T_ method. Three independent RNA extractions were performed for biological replication.

### 2.5 Western blot analysis of galectin expression

To generate protein lysates from *B. malayi* tissues, 25 adult female, 45 adult male, 300 L3 and 4x10^6^ microfilariae were homogenized with glass beads in RIPA buffer (Thermo Fischer Scientific) with Halt Protease Inhibitors (Thermo Fisher Scientific) using a disruptor genie (Scientific Industries) at 2500 rpm for 2 min followed by an incubation on ice for 2 min; a process repeated five times. Lysates were centrifuged at 14,700 x *g* for 30 min at 4°C. Protein samples were also generated from parasite extracellular vesicles (EVs). 165 adult females, 200 adult males, 2,000 L3 and 6.5x10^6^ microfilariae were cultured for 48 h and EVs isolated from spent culture media as previously described (21,23,24). Briefly, media was filtered through 0.2 μm PVDF filtered syringes (GE Healthcare) and centrifuged at 120,000 x *g* for 90 min at 4°C. The supernatant was decanted but retained, leaving approximately 1.5 ml media to ensure that the EV pellet was not disrupted. The retained media and pellet were filtered through a PVDF 0.2 μm syringe filter and centrifuged at 186,000 x *g* for a further 2 h at 4°C. The remaining supernatant was again decanted and added to previously collected supernatant leaving an isolated EV pellet. Protein was extracted from EVs by incubation in three times the volume of RIPA buffer and Halt Protease Inhibitors (Thermo Fisher Scientific) on ice for 30 min, with vortexing samples every 10 min. Following, samples were sonicated 30 s at 50% pulse then incubated on ice for 15 min; this process was repeated twice. Total protein was quantified from lysate, EVs or supernatant using the Pierce BCA Protein assay kit (Thermo Fisher Scientific) according to manufacturer’s instructions. Three µg total protein from lysate, 8 µg total protein from EVs and 480 ng of total protein from supernatant were used in subsequent western blots.

A monoclonal antibody raised against Bma-LEC-2 was provided by the Budge Lab at Washington University (St. Louis, MO) (52). A western blot of serial dilutions of rBma-LEC-2 was used to determine the sensitivity of this antibody to the recombinant protein. Proteins were resolved by SDS-PAGE using a 12% mini-PROTEAN TGX Gel (Biorad Laboratories, Hercules, CA) and transferred to a 0.2 µM nitrocellulose membrane using a Trans-Blot Turbo Mini Transfer System (Biorad Laboratories). Membranes were blocked with a blocking buffer of 5% non-fat milk powder (Cell Signaling Technology, Danvers, MA) in phosphate buffered saline with 0.05% tween-20 (PBS-T) followed by an overnight incubation at 4°C with primary antibody diluted 1:2000 in blocking buffer. The membrane was washed five times, 5 min each, with PBS-T followed by a 2 h incubation on a shaker at room temperature with goat anti-rabbit IgG-HRP (1:5000) (Thermo Fisher Scientific) in blocking buffer. The membrane was washed twice with PBS-T followed by incubation with Amersham ECL Western Blotting Detection Reagents (Cytiva). Western blots were analyzed on a Biorad Chemidoc Imaging System (Biorad Laboratories). Some SDS-PAGE gels were stained with Coomassie Brilliant Blue (Sigma-Aldrich) and imaged with a Gel Logic 112 Imaging System (Kodak, Rochester, NY).

### 2.6 CD4^+^ T cell Isolation and Apoptosis Assay

Primary naïve CD4^+^ T cells were isolated from the spleen, lymph nodes and thymus of 6–8-week-old male and female C57BL/6 WT mice (Jackson Laboratories, Bar Harbor, ME). Naïve CD4^+^ T cells were isolated using the Mojosort Mouse CD4 Naïve T Cell Isolation Kit (Biolegend, San Diego, CA) using LS columns on a QuadroMACS magnet (Miltenyi Biotec, Bergisch Gladbach, North Rhine-Westphalia, Germany). 5x10^5^ cells were plated per well of a 48-well plate coated with CD3 (5 µg/ml) (Biolegend) and cultured in regular T cell media (RPMI 1640 supplemented with 10% heat inactivated FBS (Thermo Fisher Scientific), 1% Penicillin (10,000 U/ml) (Thermo Fisher Scientific), 1% Streptomycin (10,000 ug/ml) (Thermo Fisher Scientific), 1% 100x MEM NEAA (Thermo Fisher Scientific), 1% 200 mM L-glutamine (Thermo Fisher Scientific), 55 µM β-mercaptoethanol (Thermo Fisher Scientific), CD28 (1 µg/ml) (Biolegend, San Diego, CA), and IL-2 (20 ng/ml) (Biolegend)) for naïve T cells (Th0) or polarized to Th1 phenotype by culturing in T cell media additionally supplemented with IL-12 (20 ng/mL) and anti-IL-4 (10 µg/ml) (Biolegend). After 48 h, cells were transferred to a non-CD3 coated plate and incubated for an additional three days before activation with Cell Activation cocktail with Brefeldin A (Biolegend) for 6 h and collection. To analyze apoptosis, naïve and Th1 cells were counted and plated in a 48-well plate at 5x10^5^ cells per well. Cells were incubated overnight to acclimate and then treated with either 1 µM Staurosporine (Sigma-Aldrich), dPBS (Thermo Fisher Scientific), 1 µM rBma-LEC-1 or 1 µM rBma-LEC-2. 24 h post-treatment, cells were collected and stained using the eBioscience Annexin V Apoptosis Detection Kit APC (Thermo Fisher Scientific) according to manufacturer’s instructions. Tim-3 was neutralized using *InVivo*MAb anti-mouse Tim-3 antibodies (10µg/mL) (Bio X Cell, Lebanon, NH) for 1 h prior to rBma-LEC-1 or rBma-LEC-2 treatment.

### 2.7 Macrophage Polarization Assay

The *Homo sapiens* monocyte derived THP-1 cell line (ATCC, Manassas, VA) was maintained in RPMI 1640 supplemented with 10% heat inactivated FBS, 1% 1M HEPES, 1% 100mM Sodium Pyruvate, 1% Penicillin (10,000 U/ml) 1% Streptomycin (10,000 ug/ml), 1 µg/ml Amphotericin B, 1.5 g/L Sodium Bicarbonate, and 4.5 g/L glucose (all Thermo Fisher Scientific) at 37°C with 5% CO_2_. THP-1 monocytes were seeded at 5x10^5^ per well of a 12-well plate (Thermo Fischer Scientific) and transitioned to macrophages by treatment with 80 nM Phorbol 12-myristate 13-acetate (PMA) for 2 h. Following a 24 h resting period, media was changed and cells were treated with either dPBS (control) (Thermo Fisher Scientific), 50 ng/ml IFNγ (Biolegend) + 100 ng/ml Lipopolysaccharide (LPS) (Sigma-Aldrich) for polarization to M1 phenotype, 20 ng/ml IL-4 + 20 ng/ml IL-13 (Biolegend) for polarization to M2, 0.5 µM rBma-LEC-1, 0.5 µM rBma-LEC-2 or a combination of cytokines and recombinant galectin. To investigate whether Bma-LEC-1 and Bma-LEC-2 were functioning through the Tim-3 receptor, Tim-3 was either neutralized by anti-human Tim-3 antibodies (10 µg/mL) (Biolegend) for 1 h prior to treatment or through knockdown via duplexed siRNA (Integrated DNA Technologies, Coralville, IA) (Supplemental Materials 1) treatment for 24 h prior to treatment. Cells were incubated for 72 h and then collected in Trizol for analysis of M1 and M2 markers using RT-qPCR as previously described except the High-Capacity cDNA Reverse Transcription Kit (Thermo Fisher Scientific) was used according to manufacturer’s instructions for cDNA synthesis. Expression of *TNFα, CXCL10, CD80, MCP-1, IL-10, CCL13 and CCL22* were normalized to the housekeeping gene *RPL37A* (Supplemental Materials 1).

### 2.8 Enzyme-linked Immunosorbent Assay (ELISA)

Supernatants from treated macrophage samples were collected for quantification of IL-10. An ELISA MAX Deluxe Set Human IL-10 kit (Biolegend) was used according to manufacturer’s instructions. Briefly, IL-10 capture antibody was coated to a 96-well plate and incubated at 4°C overnight. The next day the plate was washed four times with wash buffer (dPBS + 0.05% Tween-20) (Thermo Fisher Scientific) then blocked for 1 h at room temperature with assay diluent. Following, the plate was washed four times with wash buffer and 100 µl of standard or sample was added per well and incubated at room temperature for 2 h. After, the plate was washed a further four times and IL-10 detection antibody was added and incubated for 1 h. Following, the plate was washed four times and Avidin-HRP added for 30 min. After the plate was washed five times and a 3,3′,5,5′-Tetramethylbenzidine solution (Biolegend) was prepared and incubated for 30 min. Immediately after, stop solution was added and the plate was read on a SpectraMax M2e plate reader (Molecular Devices, San Jose, CA, USA).

### 2.9 Statistical Analysis

Gene expression assays were analyzed using a Two-way ANOVA with a Tukey multiple comparisons test. Apoptosis assays were analyzed using a Two-Way ANOVA Mixed Effects Analysis. Multiple comparisons were analyzed with a Šidák statistical hypothesis testing method. Macrophage polarization qPCR assays were analyzed using an Ordinary One-Way ANOVA and a Dunnett multiple comparison test. ELISA data was analyzed using a Mixed Effects Analysis model with a Holm-Šidák multiple comparison test. For all significance testing p-values < 0.05 was considered significant. All ANOVAs were completed using Graphpad prism 9.3.1 (Graphpad Software, San Diego, CA).

## 3. Results

### 3.1 Bma-LEC-1 and -2 expression and secretion are sex- and life stage-specific

We identified Bma-LEC-1 and Bma-LEC-2 as amongst the most abundant cargo proteins in EVs isolated from mature adult female *B. malayi* spent culture media but absent from EVs secreted by adult male or L3 stage parasites (21, 23). Here we used RT-qPCR to further examine the expression profile of both Bma-LEC-1 and Bma-LEC-2 across life stages of *B. malayi*. We found that, consistent with our proteomic analysis of secreted EVs, *Bma-LEC-1* and *Bma-LEC-2* expression was highest in adult female *B. malayi. Bma-LEC-1* was expressed approximately 24 fold higher in adult females than both adult males (p = 0.0345, N = 3) and microfilariae (p = 0.0338, N = 3) and 20 fold higher than in infective L3 (p = 0.0357, N=3). Adult males, infective L3 and microfilariae all had similar levels of *Bma-LEC-1* expression. Expression of *Bma-LEC-2* was also highest in adult female stages and although not statistically significant due to higher variability, adult females expressed *Bma-LEC-2* approximately five fold higher than adult males (p = 0.6286, N = 3), seven fold higher than microfilariae (p = 0.5746, N =3), and nineteen fold higher than L3 (p = 0.4973, N = 3) (Fig 1A).

**Figure 1.**
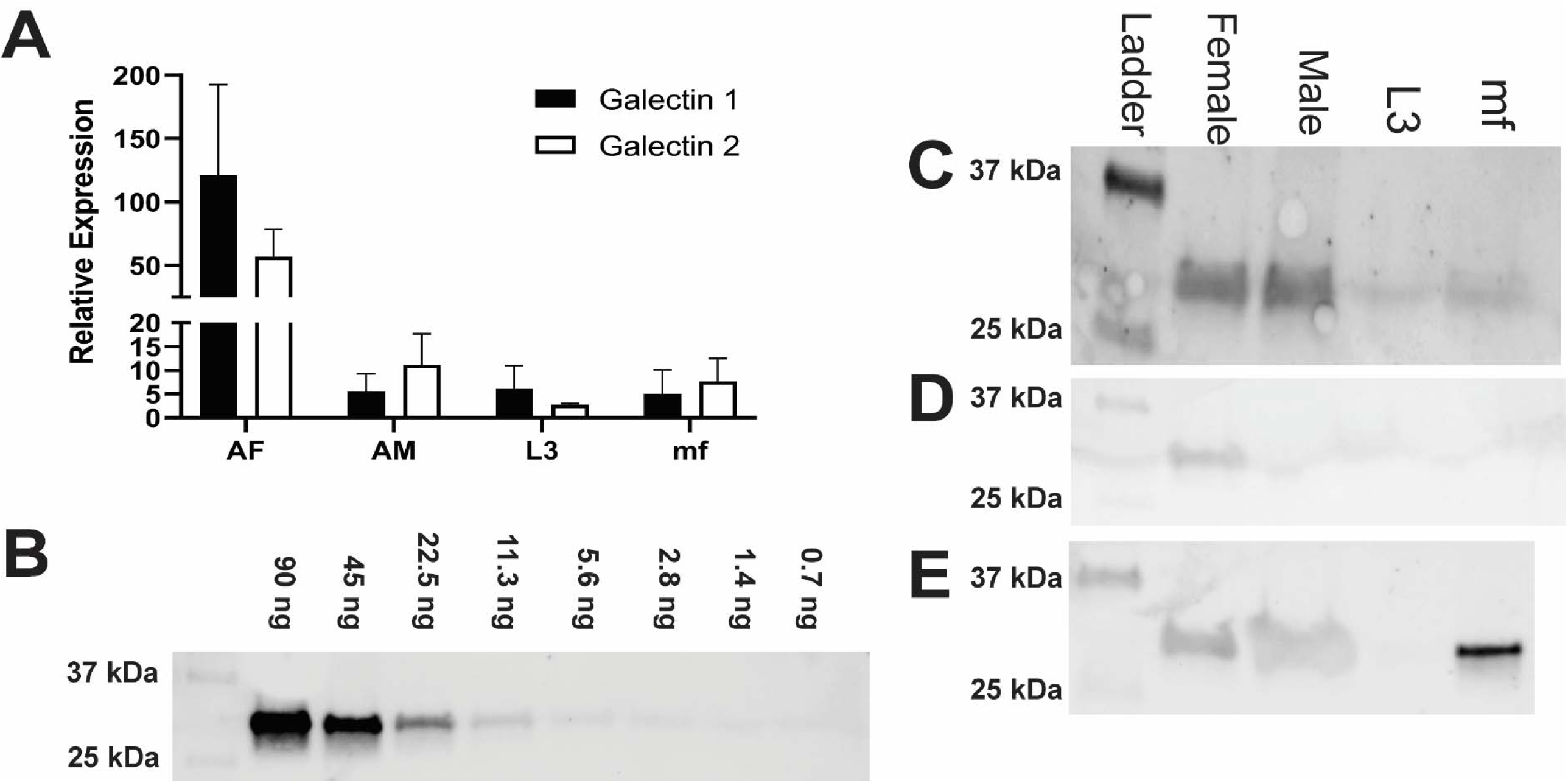
Bma-LEC-1 and Bma-LEC-2 expression and secretion are sex- and life stage-specific. (A) Bma-LEC-1 and Bma-LEC-2 are most highly expressed in adult female stage parasites. (B) A monoclonal antibody raised against Bma-LEC-2 (52) can detect recombinant Bma-LEC-2 (approximately 32 kDa) at quantities as low as 3 ng. (C) Using this antibody, parasite galectin reactivity was predominantly detected in whole worm lysate of adult female parasites and to a lesser extent in adult males and microfilariae. (D) Parasites also secrete galectins into the extracellular milieu and galectin reactivity was detected in isolated extracellular vesicles (EVs) of adult female parasites and (E) in EV-depleted spent culture media of microfilariae and to a lesser extent adult female and male parasites.

Bma-LEC-1 and Bma-LEC-2 do not contain signal peptide sequences such that their secretion into the host would require a non-canonical pathway; their identification in EVs is consistent to this. To further explore this mechanism of parasite galectin secretion, we used western blot to examine galectin expression in whole worm lysates, secreted EVs and also in EV-depleted secretory products of each life stage of *B. malayi* using an antibody raised against Bma-LEC-2 (52). It has been reported, however, that this antibody has affinity to other *B. malayi* galectins and should perhaps be viewed as a general *B. malayi* galectin antibody. Pilot experiments using recombinant Bma-LEC-2 determined that we could detect parasite galectins at levels as low as 3 ng (Fig 1B). *B. malayi* galectins were clearly identified from whole worm lysates of adult female and adult male parasites (Fig 1C). Reduced activity band was observed in whole worm lysate from microfilariae and almost none identified from L3 stage parasites. These data align well to the combined gene expression profiles of Bma-LEC-1 and Bma-LEC-2 that indicated highest galectin expression in adult stages. Galectins were also identified in protein isolated from adult female EVs, but not from EVs of any other life stage (Fig 1D). Again, this corroborates previous proteomic analyses of *B. malayi* EVs by our group and others and indicates that although galectins are being expressed endogenously, EV secretion is only by the adult female stage and hints more broadly that cargo selection and loading into EVs may be both selective and sex specific. Protein was concentrated from EV-depleted secretory products of each life stage representing proteins that are freely secreted from the parasites. Galectins were again identified in the freely secreted products of adult females and adult male parasites (Fig 1E) but surprisingly, strong reactivity was also identified in microfilariae EV-depleted secretions. Given that Bma-LEC-1 and -2 lack signal sequences, our observation of galectins in freely secreted proteins could result from EVs rupturing during sample preparation or from non-canonical secretory pathways other than EVs contributing to galectin release. The very strong reactivity in microfilariae EV-depleted products might also support this.

### 3.2 *B. malayi* galectins are related to other tandem-repeat type galectins

Clustal Omega software (53), with default settings, was used for multiple sequence alignment of selected mammalian and invertebrate galectins and c-type lectins and color coded according to Clustalx parameters. The protein sequence of *B. malayi* galectins has high similarity to other galectins found within filarial species (52), however, knowledge on the functions of this filarial nematode galectin family is lacking. A phylogenetic analysis of Bma-LEC-1 and Bma-LEC-2 was conducted to aid in the identification of potential functions. Bma-LEC-1 and Bma-LEC-2 were compared to galectins from *Caenorhabditis elegans, Drosophila melanogaster, Homo sapiens* and *Mus musculus*. A C-type lectin from each species was included to explore the relationship between Bma-LEC-1, Bma-LEC-2 and other carbohydrate binding protein types. Of the sequences used in our analysis, we found that Bma-LEC-1 and Bma-LEC-2 are most closely related to a *C. elegans* tandem-repeat type galectin, but also clustered with the tandem-repeat type galectins of humans and mice (Fig. 2A). Focusing our analysis on these tandem-repeat type galectins, we identified that Bma-LEC-1 and Bma-LEC-2 are most similar to galectin-9 proteins (Fig. 2B). Further, the similarity of the two characteristic carbohydrate recognition domains (CRDs) of the various galectins was evaluated through multiple sequence alignment (Fig. 2C-D). We found that within these critical CRDs, Bma-LEC-1 had the most similarity to *C. elegans* LEC-1 with 97% cover (percent of the query sequence found within the target sequence) and 83% identity (how similar the sequences are). Bma-LEC-1 also had high similarity to human/mouse galectin-9 CRDs with 94% cover and 60% identity and mouse galectin-4 with 92% cover and 52% identity. We also found that Bma-LEC-2 had highest similarity to *C. elegans* LEC-1 with 99% cover and 71% identity. Bma-LEC-2 also had high similarity to human/mouse galectin-9 with 92% cover and 59% identity and to human/mouse galectin-4 with 92% cover and 58% identity. These data suggest that Bma-LEC-1 and -2 are tandem-repeat type galectins and, compared to the more well understood mammalian galectins, are most similar to mammalian galectin-9.

**Figure 2.**
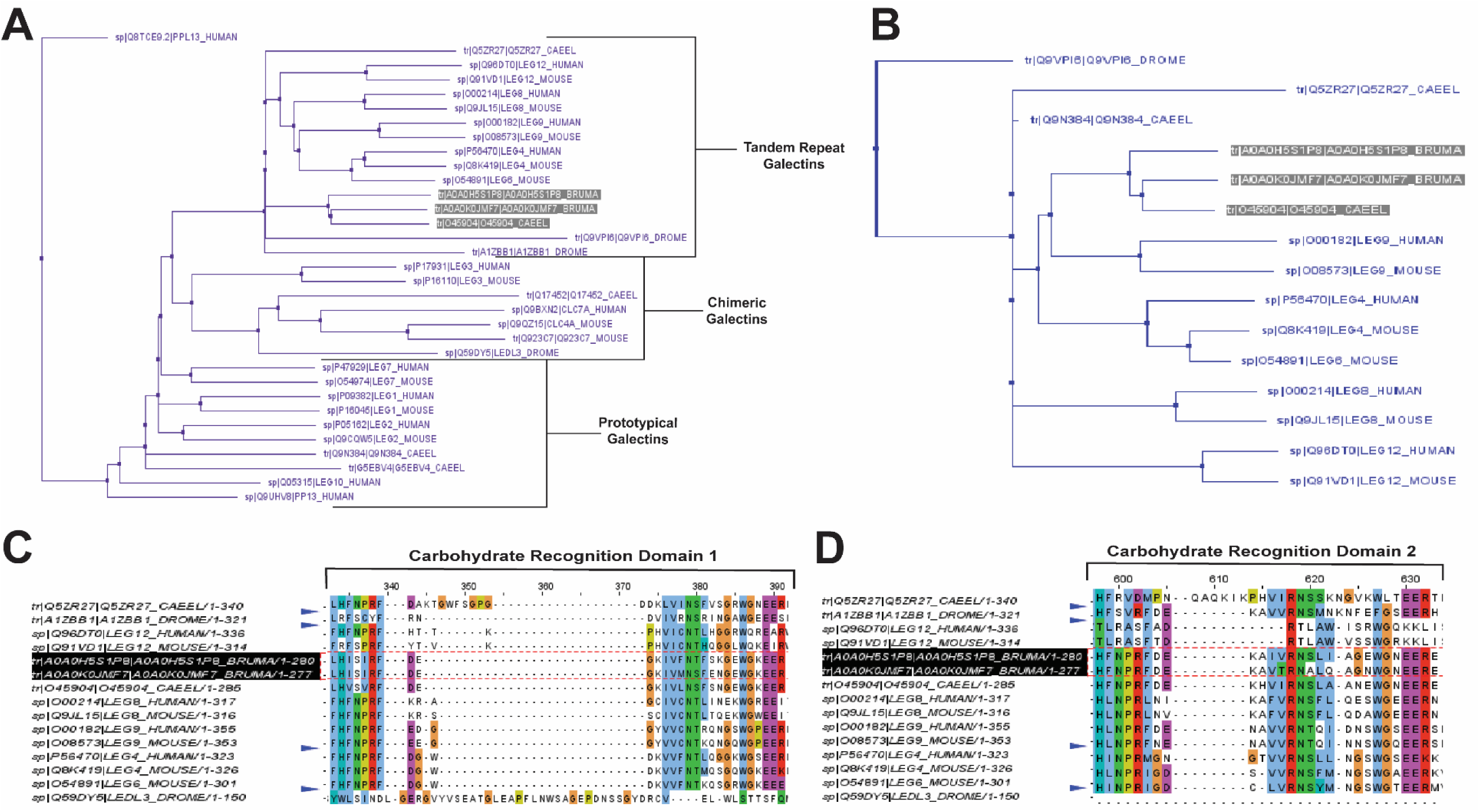
Bma-LEC-1 and -2 are similar to other tandem-repeat type galectins. (A) Bma-LEC-1 and Bma-LEC-2 cluster with other tandem-repeat type galectins from diverse species. (B) Within the tandem-repeat galectins compared, Bma-LEC-1 and Bma-LEC-2 are most closely related to *C. elegans* LEC-1 and human/mouse galectin-9. (C) Multiple sequence alignment of the two carbohydrate recognition domains characteristic of galectins revealed that Bma-LEC-1 had 83% identity to *C. elegans* LEC-1 and 60% identity to human/mouse galectin-9, while Bma-LEC-2 had 71% identity to *C. elegans* LEC-1 and 59% and 58% identity to human/mouse galectin-9, respectively. Jalview (version 2.11.1.4) was used to create phylogenetic trees and multiple sequence alignment figures.

### 3.3 rBma-LEC-1 and rBma-LEC-2 are functional homologs of mammalian galectin-9

Following cloning (Fig. 3A) and expression, recombinant protein was isolated from Sf21 cell lysates and initially purified on a Ni-NTA Resin column to bind a C-terminal His-tag. SDS-PAGE gel electrophoresis stained with Coomassie Blue revealed two distinct proteins of approximately 32 kDA in size from the elution fraction of our Ni-NTA column (Fig. 3B). Size exclusion chromatography on a FPLC system was used as a secondary additional technique to remove non-specific proteins and ensure the purity of our recombinant galectins for downstream analysis. Only FPLC fractions that contained no debris or other non-specific proteins and contained the protein of interest were used and further concentrated (Fig. 3C). Concentrated, purified proteins were confirmed by western blot using an HRP conjugated 6x-His Tag monoclonal antibody (Fig. 3D).

**Figure 3.**
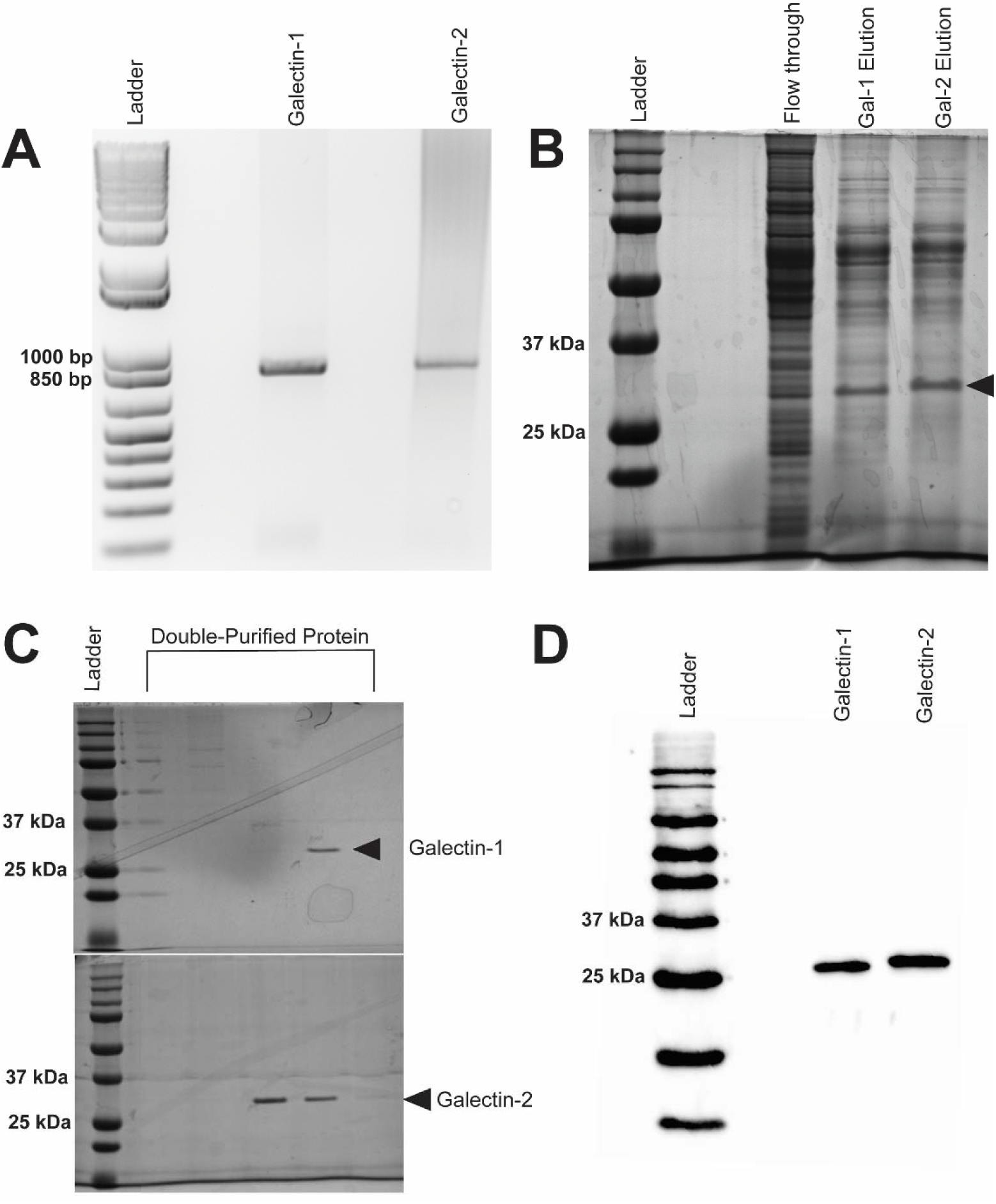
Expression of recombinant Bma-LEC-1 and Bma-LEC-2 in Sf21 cells. (A) DNA gel electrophoresis of *Bma-LEC-1* and *Bma-LEC-2* PCR amplicons from *B. malayi* adult female cDNA. (B) SDS-PAGE gel stained with Coomassie Blue showing elution of rBma-LEC-1 and rBma-LEC-2 expressing Sf21 cell lysates from initial nickel NTA resin purification and (C) after double purification using a FPLC system. Only clean FPLC fractions containing the protein of interest and no other debris or non-specific proteins were used for downstream assays. (D) Final confirmation of concentrated, double-purified rBma-LEC-1 and rBma-LEC-2 was conducted via an anti-6x His Tag western blot. Black arrowhead referencing tagged recombinant galectins of the expected 32 kDa size.

Hemagglutination (HA) and hemagglutination inhibition (HI) assays were used to obtain semi-quantitative data on the carbohydrate binding and specificities of rBma-LEC-1 and rBma-LEC-2. An HA assay was used to determine if rBma-LEC-1 and rBma-LEC-2 were expressed accurately and had produced functional proteins. Initial experiments confirmed that rBma-LEC-1 and rBma-LEC-2 were functional and capable of binding carbohydrates on the surface of rabbit erythrocytes at amounts as low as 1.5 ng and 7 ng respectively (Supplemental Materials 2). An HI assay was conducted to determine whether rBma-LEC-1 and rBma-LEC-2 had similar carbohydrate binding profiles to mammalian galectin-9, with a panel of common carbohydrates including 1 M galactose, 900 mM N-acetylgalactosamine (GalNAc), 1 M lactose, 32.6 mM N-acetyllactosamine (LacNAc), 2 M mannose, and 2 M glucose chosen for initial investigation.

The ability of rBma-LEC-1, rBma-LEC-2, human galectin-9 and mouse galectin-9 to bind RBCs were all inhibited by the presence of galactose, GalNAc, lactose and LacNAc, confirming carbohydrate binding by these galectins. This was as expected since binding these carbohydrates are properties common to lectin-type proteins (48). In addition, the ability of rBma-LEC-1, rBma-LEC-2, human galectin-9 and mouse galectin-9 to bind RBCs was not inhibited by glucose or mannose (Fig 4A). Again, this was expected as these sugars are not common binding targets of tandem-repeat galectins (54, 55). The affinities of these galectins to galactose, GalNAc, lactose, and LacNAc were further investigated by determining the minimal inhibitory concentration (MIC) of the carbohydrate solution that could still inhibit hemagglutination. The MIC of galactose for human galectin-9 and mouse galectin-9 was determined to be 125 mM and 62.5 mM respectively. rBma-LEC-2 was similar with a MIC of 125 mM but rBma-LEC-1 bound galactose with greater sensitivity (MIC < 488 µM). The MIC of GalNAc for both human and mouse galectin-9 was 112.5 mM, while rBMA-LEC-2 had a MIC of 450 mM; again, rBma-LEC-1 had the highest sensitivity for GalNAc binding with a MIC of < 440 µM. There was more variation in the galectin’s ability to bind to lactose as compared to other carbohydrates. Human galectin-9 and rBma-LEC-1 had a MIC for lactose of < 488 µM, while mouse galectin-9 and rBma-LEC-2 had MICs of 4 mM and 15.6 mM respectively. All galectins tested had the lowest MIC and thus highest sensitivity to LacNAc with human galectin-9 and rBma-LEC-1 having a MIC of < 255 µM and mouse galectin-9 and rBma-LEC-2 a MIC of 2 mM (Fig 4B-C) the lowest MIC of any of the carbohydrates tested.

**Figure 4.**
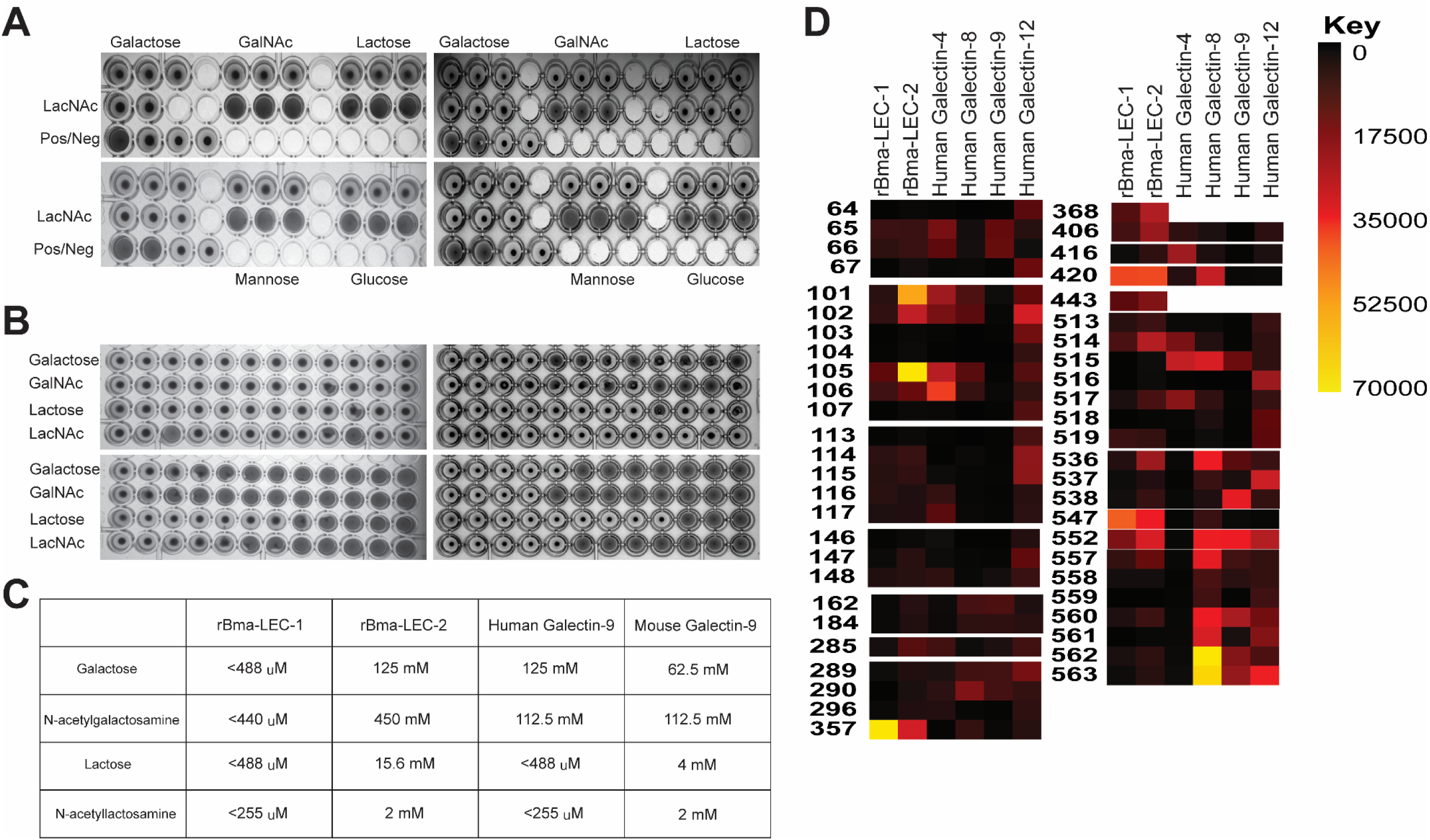
rBma-LEC-1/LEC-2 are functional homologs of mammalian galectin-9. (A) Hemagglutination inhibition (HI) assay was used to determine the carbohydrate binding specificities of rBma-LEC-1 and rBma-LEC-2. A panel of common carbohydrates was used to test if the recombinant proteins were functional galectins. A solid colored well indicates that the galectin present is not binding to the carbohydrate of interest but is binding the erythrocytes instead forming a lattice of erythrocytes on the bottom of the well. A well with a dot in the center indicates that the galectin present is binding the carbohydrate of interest not the erythrocytes allowing them to sediment at the bottom of the well. Both recombinant proteins (top and bottom left) were capable of binding galactose, GlcNAc, lactose, and LacNAc, each known galectin substrates. A similar sugar-binding profile was observed with human (top right) and mouse galectin-9 (bottom right). (B) The binding sensitivity of galectins were determined using a minimal inhibitory concentration (MIC) of each carbohydrate solution. Wells with a dot in the center indicate that the galectin is able to bind that concentration of the carbohydrate of interest. The MIC is determined at the first well where the galectin is no longer able to bind that concentration of carbohydrate as indicated by a solid colored well. rBma-LEC-1 (top left) had the highest affinity to all of the carbohydrate solutions tested and all galectins had the highest affinity to LacNAc. (C) A glycan binding array was used to compare rBma-LEC-1 and rBma- LEC-2 glycan binding profiles with those of human galectin -4, -8, -9 and -12 using a substrate of 584 glycan moieties. The glycan binding profiles of the parasite galectins are most similar to human galectin-8 and galectin-9. Glycan binding arrays are quantified by relative fluorescence units (RFU). An abbreviated version of this array data is presented with a full analysis found in Supplemental Materials 3. A corresponding key of glycan names associated with the numerical IDs is provided in Supplemental Materials 4.

HA and HI assays are robust but not high throughput so a glycan-binding array was performed to comprehensively profile the binding specificities of rBma-LEC-1 and rBma-LEC-2 compared to mammalian, tandem-repeat type galectins. Glycan binding array data was compared to archived data of human galectin-4, galectin-8, galectin-9 and galectin-12 available at the National Center for Functional Glycomics (https://ncfg.hms.harvard.edu/). rBma-LEC-1 and rBma-LEC-2 had the most similar glycan binding patterns to human galectin-8 and human galectin-9 (Fig. 4D).

These parasite galectins shared mammalian galectin-8 and -9 ability to bind to Gala1-2Galb-CH2CH2CH2NH2, Gala1-3(Fuca1-2)Galb1-3GlcNAcb-CH2CH2NH2, Gala1-3(Fuca1-2)Galb1-4(Fuca1-3)GlcNAcb-CH2CH2CH2NH2, Galb1-4(Fuca1-3)GlcNAcb1-2Mana1-6(Galb1-4(Fuca1-3)GlcNAcb1-2Mana1-3)Manb1-4GlcNAcb1-4(Fuca1-6)GlcNAcb-NST, GlcNAcb1-3Galb1-4GlcNAcb1-6(GlcNAcb1-3)Galb1-4GlcNAc-CH2CH2NH2, and Galb1-3GlcNAcb1-3Galb1-4GlcNAcb1-2Mana1-6(Galb1-3GlcNAcb1-3Galb1-4GlcNAcb1-2Mana1-3)Manb1-4GlcNAcb1-4GlcNAc-VANK. This strong glycan binding congruency indicates that rBma-LEC-1 and rBma-LEC-2 may have the ability to phenocopy human galectin-8 and galectin-9 if secreted by the parasite into similar contexts as the endogenous host galectins. The full glycan binding array data for rBma-LEC-1 and rBma-LEC-2 can be found in Supplemental Materials 3 with a key provided in Supplemental Materials 4. Overall, rBma-LEC-1 was able to bind to more glycans on the array than rBma-LEC-2. Both recombinant galectins had a particularly high affinity for O-glycan and N-glycan motifs that form blood group B antigens. More specifically, out of all the glycans assayed, rBMA-LEC-1 had the highest affinity to alpha-galactose on a biantennary N-glycan, to blood group B on a biantennary N-glycan, and to blood group B on multiple O-glycan and N-glycan motifs while rBma-LEC-2 had the highest affinity to blood group B, to blood group B on biantennary N-glycans, to alpha-galactose on biantennary N-glycans, and to various O-glycan and N-glycan motifs. Structures of the glycans that rBma-LEC-1 and rBma-LEC-2 had the highest affinity for can be found in Figure 5.

**Figure 5.**
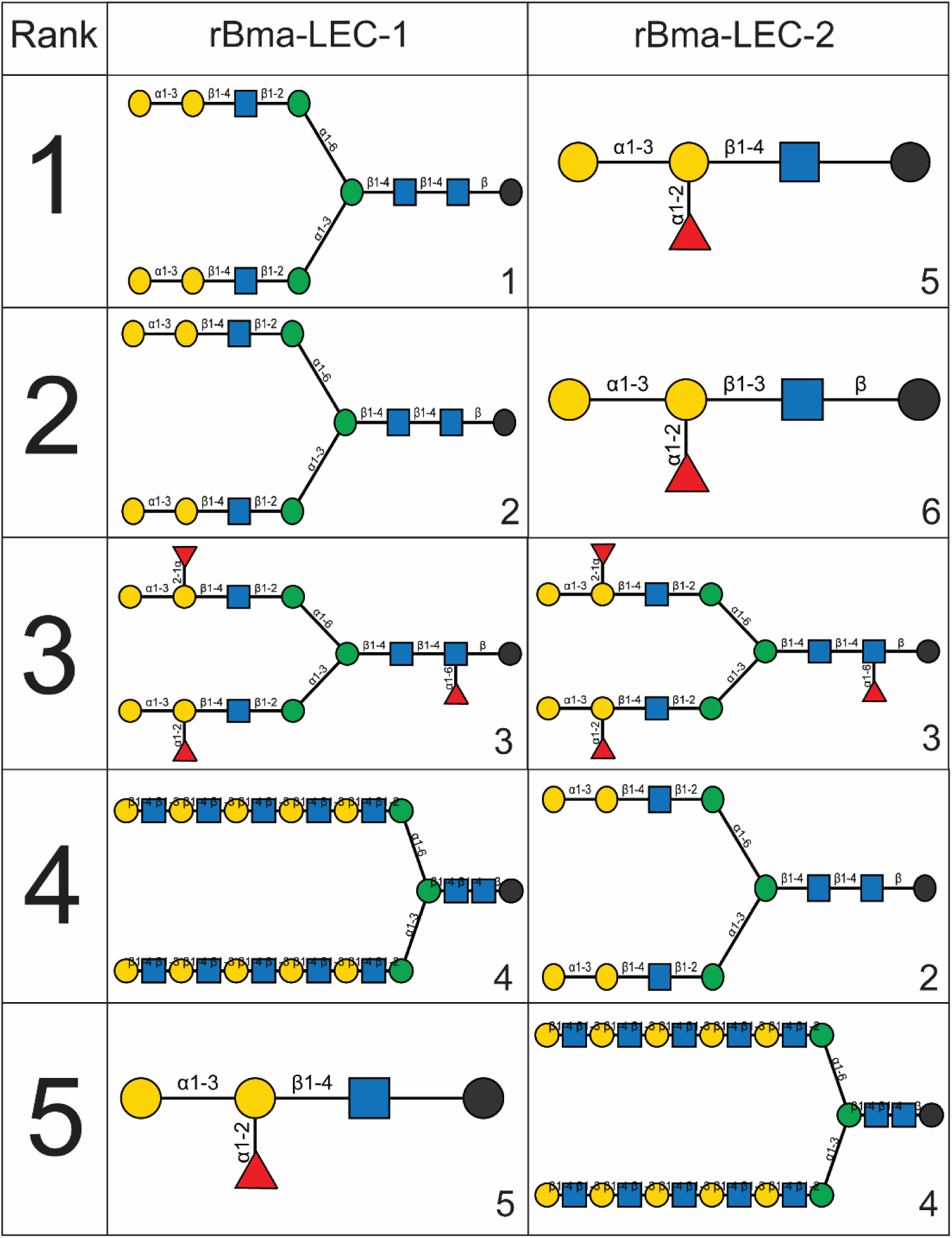
Structures of rBma-LEC-1/LEC-2 highest affinity glycans. The binding affinities of rBma-LEC-1 and rBma-LEC-2 were investigated with a glycan binding array. rBma-LEC-1 had high affinity for alpha-galactose on a biantennary N-glycan, for blood group B on a biantennary N-glycan, and for blood group B on multiple O-glycan and N-glycan motifs. rBma-LEC-2 had high affinity for blood group B, for blood group B on biantennary N-glycans, for alpha-galactose on biantennary N-glycans, and for various O-glycan and N-glycan motifs. This table shows the structures of the top five highest binding glycans. 1: Gala1-3Galb1-4GlcNAcb1-2Mana1-6(Gala1-3Galb1-4GlcNAcb1-2Mana1-3)Manb1-4GlcNAcb1-4GlcNAcb-GENR, 2: Gala1-3Galb1-4GlcNAcb1-2Mana1-6(Gala1-3Galb1-4GlcNAcb1-2Mana1-3)Manb1-4GlcNAcb1-4GlcNAc-KVANKT, 3: Gala1-3(Fuca1-2)Galb1-4GlcNAcb1-2Mana1-6(Gala1-3(Fuca1-2)Galb1-4GlcNAcb1-2Mana1-3)Manb1-4GlcNAcb1-4(Fuca1-6)GlcNAcb-NST, 4: Galb1-4GlcNAcb1-3Galb1-4GlcNAcb1-3Galb1-4GlcNAcb1-3Galb1-4GlcNAcb1-3Galb1-4GlcNAcb1-2Mana1-6(Galb1-4GlcNAcb1-3Galb1-4GlcNAcb1-3Galb1-4GlcNAcb1-3Galb1-4GlcNAcb1-3Galb1-4GlcNAcb1-2Mana1-3)Manb1-4GlcNAcb1-4GlcNAcb-VANK, 5: Gala1-3(Fuca1-2)Galb1-4GlcNAc-CH2CH2NH2, 6: Gala1-3(Fuca1-2)Galb1-3GlcNAcb-CH2CH2NH2.

### 3.4 rBma-LEC-2, but not rBma-LEC-1, induces immunomodulatory phenotypes seen in chronic filarial infection

Mammalian galectin-9 is a known immunomodulatory molecule and promotes suppressive or regulatory phenotypes in both lymphoid and myeloid immune cells. For example, it has been shown that mammalian galectin-9 can induce apoptosis in CD4^+^ Th1 cells (56–59) and promote polarization of macrophages to an alternatively activated or M2 phenotype (59–62). These phenotypes, among others elicited by galectin-9, are consistent with the modifications of the initial type 2 immune response seen during chronic filarial infection. Due to the similar glycan binding profiles of Bma-LEC-1, Bma-LEC-2 and mammalian galectin-9 we wanted to investigate whether parasite-derived galectins could induce similar phenotypes. To investigate whether rBma-LEC-1 and rBma-LEC-2 could induce apoptosis in Th1 cells, primary naïve CD4^+^ T cells were isolated from the spleens, lymph nodes and thymus of C57BL/6 mice and a subset polarized to Th1. Both naïve and Th1 cells were treated with rBma-LEC-1 and rBma-LEC-2 and apoptosis was detected using Annexin V conjugation and quantified with flow cytometry. Our ability to measure apoptosis using this approach was confirmed using 1 µM staurosporine as a positive control, which increased apoptosis in naïve and Th1 cells by 49% and 44%, respectively (p < 0.0001, N = 8) compared to negative control CD4^+^ cells treated with dPBS. Treatment with 1 µM rBMA-LEC-1 significantly increased apoptosis in Th1 cells by 13% as compared to rBma-LEC-1 treated naïve T cells (p = 0.0137, N = 6), while treatment with rBma-LEC-2 significantly increased apoptosis in Th1 cells by 20% as compared to rBma-LEC-2 treated naïve T cells (p < 0.0001, N = 7). However, only rBma-LEC-2, not rBma-LEC-1, treated Th1 cells had significantly higher apoptosis than negative control Th1 cells. rBma-LEC-2 treated Th1 cells had a 15% increase (p = 0.0059, N = 7) in apoptosis as compared to negative control Th1 cells (Fig. 6A). This data support the conclusion that although rBma-LEC-1 displays an *in vitro* glycan binding profile similar to mammalian galectin-9, it does not phenocopy the reported functional effects of galectin-9 in this assay. In contrast, the similar rBma-LEC-2 does phenocopy and can selectively induce apoptosis in Th1 cells, but not naïve CD4^+^ T cells. The mechanism of apoptosis induction in Th1 cells by mammalian galectin-9 is known to involve binding T cell immunoglobulin and mucin domain-containing protein 3 (Tim-3) on the surface of CD4^+^ T cells (56). To determine if rBma-LEC-2 utilizes the same receptor to induce apoptosis we neutralized the Tim-3 receptor by treatment with anti-mouse Tim-3 antibodies. Neutralization of Tim-3 did not abrogate the effects of rBma-LEC-2 as expected (Fig. 6A), suggesting either that the antibody is not fully preventing the binding of Bma-LEC-2 or that Bma-LEC-2 is mediating its effects through alternative pathways. To explore this further we attempted to use RNAi to suppress Tim-3 receptor expression, however, we were unable to maintain isolated T cells sufficiently in culture to enable this approach.

**Figure 6.**
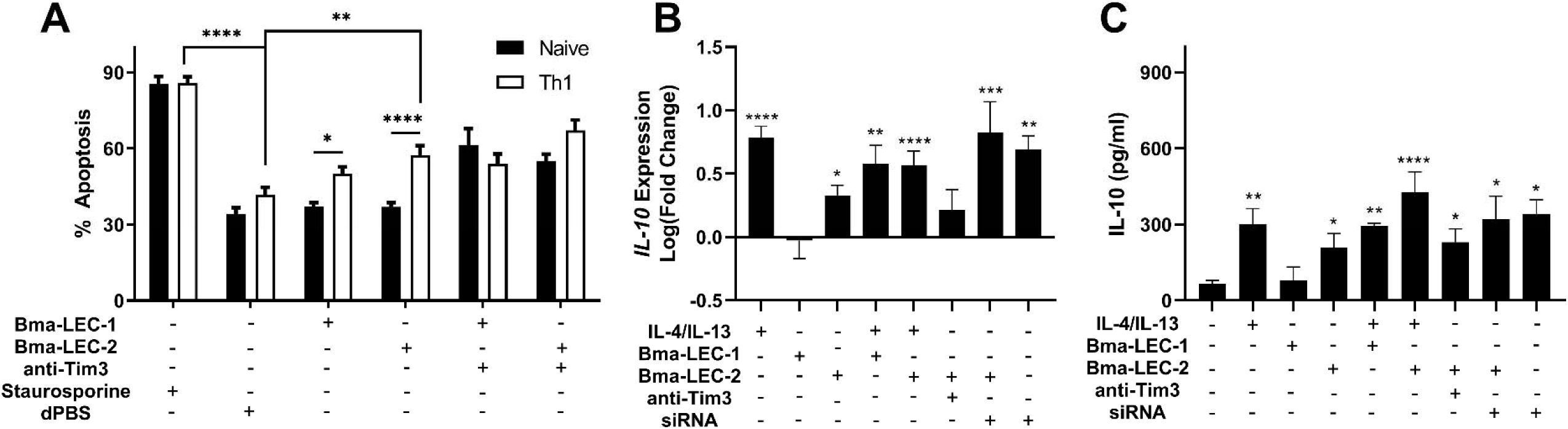
rBma-LEC-2 induces suppressive phenotypes in lymphoid and myeloid cells. (A) rBma-LEC-2, but not rBma-LEC-1, is a bioactive effector molecule that selectively induces apoptosis in Th1 cells, but not naïve T cells and (B) promotes polarization of macrophages to an alternatively activated phenotype as indicated by its increase in expression and (C) production of IL-10. These phenotypes are consistent with phenotypes induced by mammalian galectin-9. There is no synergistic effect in either inducing apoptosis of T cells or promoting polarization of macrophages when cells are treated with both human cytokines IL-4 and IL-13 and parasite-derived galectin. These effects may be mediated by T cell immunoglobulin and mucin domain-containing protein 3 (Tim-3), a known receptor for galectin-9. Suppressing the expression of Tim-3 with siRNA induced a similar increase in IL-10 as treatment of cells with Bma-LEC-2. N = 3 (minimum). Mean ± SEM, *P<0.05, **P<0.01, ***P<0.001, ****P<0.0001.

Further to inducing apoptosis in Th1 cells, galectin-9 is known to elicit alternative activation phenotypes in macrophages. To test the hypothesis that rBma-LEC-1 and rBma-LEC-2 can phenocopy galectin-9 and also drive macrophage polarization, THP-1 cells, a human monocyte cell line, were differentiated to macrophages using PMA and then treated with either rBma-LEC-1 or rBMA-LEC-2 alone. Polarization was analyzed by quantifying expression of common M1 and M2 macrophage markers including *TNFα, CXCL10, CD80, MCP-1, IL-10, CCL13 and CCL22.* Treatment with rBma-LEC-1 or rBma-LEC-2 did not promote polarization to the classically activated (M1) phenotype as indicated by the lack of expression of any of the M1 markers (Supplemental Materials 5A-C), however, treatment with rBma-LEC-2 alone did significantly increase expression of one M2 marker, *IL-10*, but not others (*MCP-1, CCL13 or* CCL22, see Supplemental Materials 5D-F). Focusing on this phenotype, we found that treatment of macrophages with rBma-LEC-1 had no effect on expression of *IL-10* (p = 0.9997, N = 10), but treatment with rBma-LEC-2 significantly increased expression by 213% (p = 0.0411, N = 13) (Fig 6B). To determine if there was any synergistic effect of parasite galectin treatment alongside traditional M2 polarization using IL-4 and IL-13 we treated macrophages with each parasite-derived galectin in combination with IL-4 and IL-13. Treatment with the combinations of rBma-LEC-1/IL-4/IL-13 and rBma-LEC-2/IL-4/IL-13 significantly increased expression of *IL-10* by 379% (p = 0.0010, N = 7) and 366% (p < 0.0001, N = 12), respectively (Fig. 6B) but did not appear synergistic as *IL-10* expression following both treatments was not significantly higher than for IL-4/IL-13 alone (p = 0.7409, N = 7 and p = 0.4602, N = 12, respectively). To evaluate whether rBma-LEC-2 driven increase in *IL-10* expression was mediated through Tim-3, the Tim-3 receptor was both neutralized using anti-human Tim-3 antibody or was knocked down by treatment with siRNA (in contrast to our T cell work, the longevity of differentiated macrophages in culture allowed this approach) (Supplemental Materials 6). Neither neutralization (p = 0.9962, N = 4) nor *Tim-3* knock down (p = 0.1112, N = 3) abrogated the effects of rBma-LEC-2 (Fig. 6B). From the antibody data, one may again conclude either that the antibody is not fully preventing the binding and therefore not fully inhibiting the effects of rBma-LEC-2 on *IL-10* expression, or that Bma-LEC-2 is not mediating its effects through Tim-3. For the siRNA approach we had expected Tim-3 knockdown to abrogate the effects of Bma-LEC-2, however, it has been shown that silencing Tim-3 on monocytes can increase *IL-10* expression (63, 64). Our data revealed a similar trend in that treatment with either siRNA + rBma-LEC-2 (p = 0.0008, N = 3) or treatment with siRNA only (p = 0.0059, N = 3) increased expression of *IL-10* as compared to naïve macrophages. In addition, neither treatment was statistically different from treatment with rBma-LEC-2 only.

In addition to analyzing *IL-10* at the transcriptional level, we also analyzed IL-10 production at the protein level using ELISA. In correlation with the gene expression data, rBma-LEC-1 did not have any effects at all on IL-10 production (0.7600, N = 4) while rBma-LEC-2 significantly increased IL-10 production by 312% (p = 0.0262, N = 9) (Fig. 6C). Again, in strong congruence with the transcriptional data, we did not observe any synergistic effects of either parasite galectin in combination with IL-4/IL-13 (Fig. 6C). Neither treatment with anti-Tim-3 antibody (p = 0.3650, N=3) nor treatment with Tim-3 siRNA (p = 0.3838, N =3) abrogated the effects of rBma-LEC-2 (Fig. 6C) but we again saw that treatment with siRNA + rBma-LEC-2 (p = 0.0262, N = 3) and treatment with siRNA only (p = 0.0223, N = 3) significantly increased production of IL-10 (Fig. 6C). Collectively, our data show that rBma-LEC-1 does not exhibit bioactivity in the *in vitro* Th1 apoptosis and macrophage differentiation assays used here but rBma-LEC-2 is bioactive, promoting apoptosis of Th1 cells but not naïve, native CD4^+^ cells and increasing expression of IL-10 in human macrophages. This activity phenocopies the reported functions of endogenous human galectin-9. The mechanism of action of rBma-LEC-2 is unclear but our data do support Tim-3 involvement. Tim-3 is an inhibitory receptor (65–69) and suppression of its expression in macrophages by siRNAs removes that brake to increase *IL-10* expression. Similarly, galectins act to bind Tim-3 and sequester its activity through their lattice-forming nature. This would also remove the Tim-3 mediated inhibition of *IL-10* expression and we see that effect in our IL-10 assays.

## 4. Discussion

The immunosuppressive phenotypes induced by Bma-LEC-2 give new insight into some of the mechanisms filarial parasites utilize to establish and maintain chronic infection. Hallmarks of chronic filarial infections include an increase in alternatively activated macrophage (3–5) and CD4^+^ CD25^+^ regulatory T cell (5–10) populations, increases in IL-4 and IL-10 (11–13), and a reduction in or a hyporesponsiveness of T cells (14–17). The outcome of these modifications is to create a state of immune tolerance where the host can maintain an active immune response, but damage to the parasite is limited. Here we have shown that protein cargo within parasite-derived EVs, specifically a parasite-derived galectin-9 homolog (Bma-LEC-2), can induce polarization of macrophages to an alternatively activated phenotype, increase *IL-10* expression in macrophages and induce apoptosis in Th1 cells.

The concept of filarial and other parasitic nematodes evolutionarily adapting to produce homologs of host molecules to aid in their survival has been documented. *B. malayi* secrete a homolog of the human cytokine macrophage migration inhibitory factor homolog 1 (MIF-1) that can phenocopy the host molecule to induce chemotaxis of human monocytes (70, 71) and increase production of IL-8 and TNFα in those cells (70). Many filarial species also secrete a homolog to host cystatins, a cysteine protease inhibitor. Onchocystatin, from *Onchocerca volvulus*, has immunomodulatory effects on human monocytes as indicated by its ability to suppress proliferation and increase production of IL-10 in human PBMCs (72). In *B. malayi*, secretion of BmCPI-2 inhibits the antigen presentation abilities of human B cell lines (73). Other host cytokine homologs secreted by *B*. *malayi* include asparaginyl-tRNA synthetase, a structural homolog to human IL-8, and tgh-2, a homolog to human TGFβ. In a T-cell transfer colitis mouse model, asparaginyl-tRNA synthetase induced IL-10 production in splenic cells and mice treated with this protein showed resolution of cellular infiltration in their colonic mucosa and induced a CD8^+^ cellular response (74). *B. malayi* tgh-2 can bind the TGFβ receptor in mink epithelial cells and activates plasminogen activator inhibitor-1 expression, a marker for TGFβ-mediated transduction (75). These phenotypes show that parasite-derived homologs of host immunoregulatory molecules can create an overall state of hyporesponsiveness in immune cells that represents a benefit to the parasite. Our findings continue to document this strategy with *B. malayi* adult females producing a homolog of mammalian galectin-9 that induce immunoregulatory phenotypes in mammalian immune cells.

The diverse and important roles that host galectins have in immunomodulation make them effective molecules for the parasite to mimic; the β-galactoside binding properties of galectins give them a wide range of cell types which they can target and secretion of parasite galectins into the host milieu may be an evolutionarily conserved strategy. While we have shown that *B. malayi* adult females secrete EVs containing a bioactive galectin that can drive immunosuppressive phenotypes, they are not the only filarial nematode to utilize galectins to potentially modulate their environment and promote survival. *Dirofilaria immits* produce a galectin-like protein that can bind plasminogen and stimulate plasmin generation by tissue plasminogen activator on canine endothelial cells, therefore activating the host fibrinolytic system (76) and stimulating the degradation of extracellular matrix via the host plasminogen/plasmin system (77). In addition to filarial nematodes, gastrointestinal nematodes have also been shown to secrete galectins. The infective L3 stage larvae of the gastrointestinal nematode *Haemonchus contortus* produces several galectin-like proteins with a at least one possessing eosinophil chemokine activity (78). Another study has shown that *H. contortus* produces Hco-GAL-1, a galectin like protein, that can increase production of IL-10 from goat monocytes, inhibit T cell proliferation and induce apoptosis in goat T cells (79), phenotypes consistent with those we have described for Bma-LEC-2. In addition, *Toxascaris leonine*, an Ascarid of canines and felines, produces a galectin-9 homolog based on its structure and CRD sequence (80). This galectin was shown to increase concentrations of TGFβ and IL-10 in pretreated dextran sulfate sodium (DSS) treated mice (81) promoting an immunosuppressive environment.

The functional role for galectin-9 driving an IL-10 regulatory environment is not just relevant at the host-helminth interface but is fundamental to host-pathogen interactions more generally. Many studies have shown that there is increased expression of galectin-9 in chronic Hepatitis C Virus (HCV) infected patients (82–84) with patients displaying an increased expansion of regulatory T cell populations and a contraction of CD4^+^ effector T cells (82, 83). An *in vitro* study examining HCV infected human hepatocytes co-cultured with CD4^+^ T cells showed that the CD4^+^ T cells had increased expression of galectin-9, TGFβ and IL-10, indicating viral infection was driving regulatory phenotypes (84). This study suggested that the virus itself is modulating host immunity through the use of galectin-9. A similar phenotype was seen in human dendritic cells (DC) infected with Dengue virus. In this context, infection with dengue virus specifically increases mRNA and protein levels of galectin-9 (85).

Our data are consistent with the effects of Bma-LEC-2 being mediated, at least in part, by the Tim-3 receptor, similar to mammalian galectin-9. However, studies on mammalian galectin-9 have indicated that mammalian galectin-9 can interact with other receptors also resulting in immunomodulatory effects. For example, mammalian galectin-9 can bind CD44 on T cells thereby inhibiting the binding of hyaluronan (HA) to CD44 (86, 87). In a study using a murine model of allergic asthma it was shown that mammalian galectin-9 can modulate CD44-dependent leukocyte recognition of the extracellular matrix leading to a reduction in airway hyperresponsiveness (86). Similarly, mammalian galectin-9 suppressed binding of HA to CD44 on melanoma and colon cancer cell lines resulted in suppression of both attachment and invasion of tumor cells by inhibiting their binding of adhesive molecules to ligands on vascular endothelium and the extracellular matrix (87). In a viral study, mammalian galectin-9 was identified as a novel binding partner of Epstein-Barr virus latent membrane protein 1 (LMP1) from nasopharyngeal carcinomas (88). These studies provide evidence that mammalian galecin-9 uses various binding partners each resulting in an immunomodulatory phenotype. Further investigation into additional potential binding partners of Bma-LEC-2 is therefore necessary to truly understand the extent of its immunomodulatory effects. The glycan binding data presented here provides a guide for the identification of these partners.

## Acknowledgments

*Brugia* life cycle stages were obtained through the NIH/NIAID Filarial Research Reagent Resource Center (FR3), morphological voucher specimens are stored at the Harold W. Manter Museum at University of Nebraska, accession numbers P2021-2032.

