## Supplemental Materials 1 for "Secreted filarial nematode galectins modulate host immune cells"

**Table 1. Primer Sequences**

| Experiment | Gene/Protein | Forward Primer | Reverse Primer |
|---|---|---|---|
| Protein Expression | | | |
| | *Bma-lec-1* (WBGene00226528) | 5'AAAGGATCCATGTCTGATCAGAG ATCATATCC3' | 5'AAAGCGGCCGCGTGAATTTGTATGCC CGTAAG3' |
| | *Bma-lec-2* (WBGene00224538) | 5'AAAAAGCTTATGGCCAATGAATATG AAACGAATTATCC-3' | 5'AAAGCGGCCGCGCATCTGGATACC GCTTAC3' |
| Galectin Expression across Life stages qPCR | | | |
| | *Bma-lec-1* | 5'-CTGCCATTGTCTTCCGTATCT-3' | 5'-TGCCTTCCCTCTCCTCATTA-3' |
| | *Bma-lec-2* | 5'- ACGCTTCCACATCAATCTACTC-3' | 5'-GCATCTGGATACCGCTTACTT-3' |
| | *B. malayi NADH Dehydrogenase Subunit 1* (NC_004298.1) | 5'-GGGTGGCACTCAGTGTCGTA-3' | 5'-ACAACGCCTGAAAAATACCAG-3' |
| Macrophage Polarization | | | |
| | *CCL13* | 5'-CCAAACTGGGCAAGGAGAT-3' | 5'GTCTTCAGGGTGTGAGCTTT-3' |
| | *CCL22* | 5'-TAGGCTCTTCATTGGCTCAG-3' | 5'-ATTACGTCCGTTACCGTCTG-3' |

| | | | |
|---|---|---|---|
| | *CD80* | 5’-ATCCTGGGCCATTACCTTAATC-3’ | 5’-CTCTCATTCCTCCTTCTCTCTCT-3’ |
| | *CXCL10* | 5’-CCATTCTGATTTGCTGCCTT-3’ | 5’-TACTAATGCTGATGCAGGTA-3’ |
| | *IL10* | 5’-TACGGCGCTGTCATCGATTT-3’ | 5’-TAGAGTCGCCACCCTGATGT-3’ |
| | *MCP1(CCL2)* | 5’-GATCTCAGTGCAGAGGCTCG-3’ | 5’-TTTGCTTGTCCAGGTGGTCC-3’ |
| | *RPL37A* | 5’-ATTGAAATCAGCCAGCACGC-3’ | 5’-AGGAACCACAGTGCCAGATCC-3’ |
| | *TNF* | 5’-TCTTCTCGAACCCCGAGTGAC-3’ | 5’-TTTGCTTGTCCAGGTGGTCC-3’ |
