## Supplemental Materials 2 for "Secreted filarial nematode galectins modulate host immune cells"

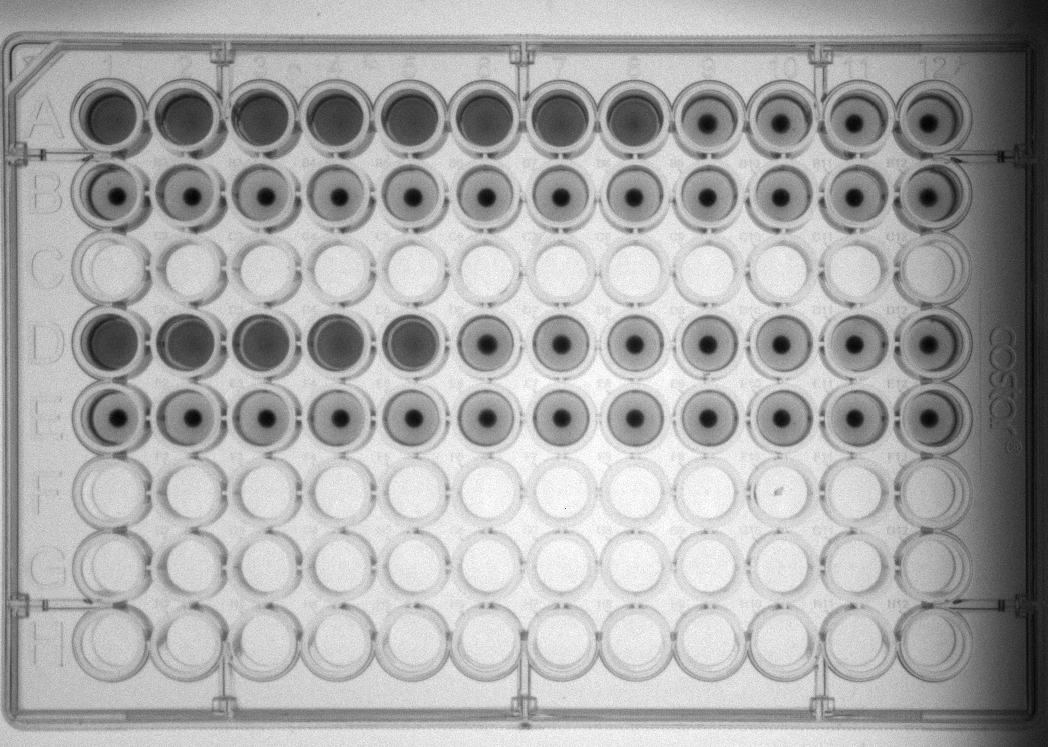

**rBma-LEC-1**

**rBma-LEC-2**

**Supplemental Materials 2: Hemagglutination Assay for rBma-LEC-1 and rBma-LEC-2**

A Hemagglutination Assay was used to determine if rBma-LEC-1 and rBma-LEC-2 were expressed appropriately and whether they were functional galectins. This was determined on their ability to bind carbohydrate moieties on the exterior of red blood cells (RBCs). Serial dilutions of rBma-LEC-1 and rBma-LEC-2 were mixed with trypsinized rabbit RBCs. It was determined that rBma-LEC-1 was capable of binding RBCs at concentrations as low as 1.5 ng while rBma-LEC-2 could bind at concentrations as low as 7 ng.
