## Supplemental Materials 5 for "Secreted filarial nematode galectins modulate host immune cells"

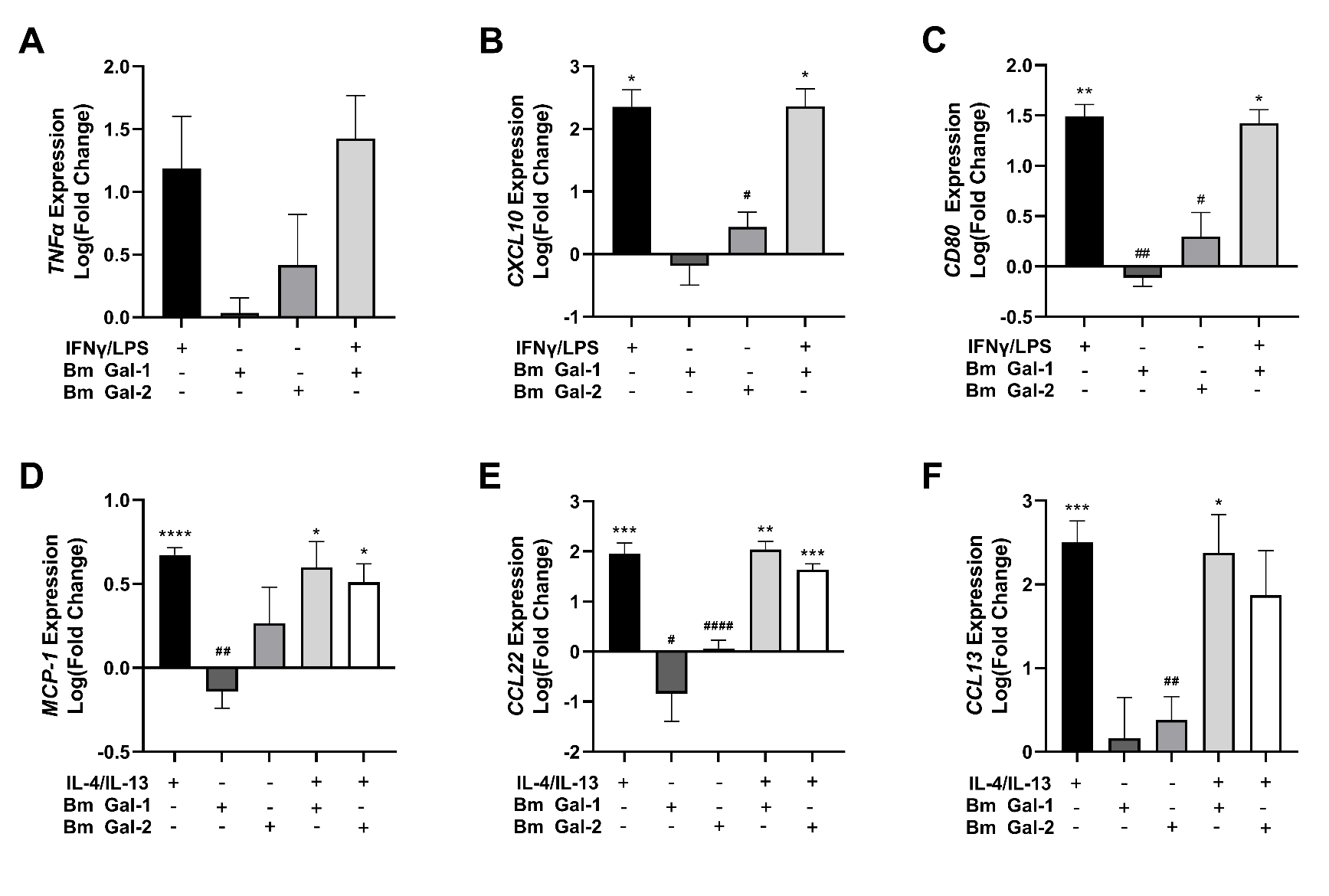

**Supplemental Materials 5. rBma-LEC-1 and rBma-LEC-2 do not increase M1 markers and some M2 markers**

rBma-LEC-1 and rBma-LEC-2 did not statistically increase any of the M1 markers tested (A-C). While rBma-LEC-2 did statistically increase IL-10, it did not statistically increase other M2 markers that were tested including *MCP-1, CCL22, and CCL13* (D-F). N = 3 (minimum). Mean ± SEM, * = Statistical significance relative to M0. *P<0.05, **P<0.01, ***P<0.001, ****P<0.0001. # = Statistical significance relative to M2. #P<0.05, ##P<0.01, ####P<0.0001.
