## Supplemental Materials 6 for "Secreted filarial nematode galectins modulate host immune cells"

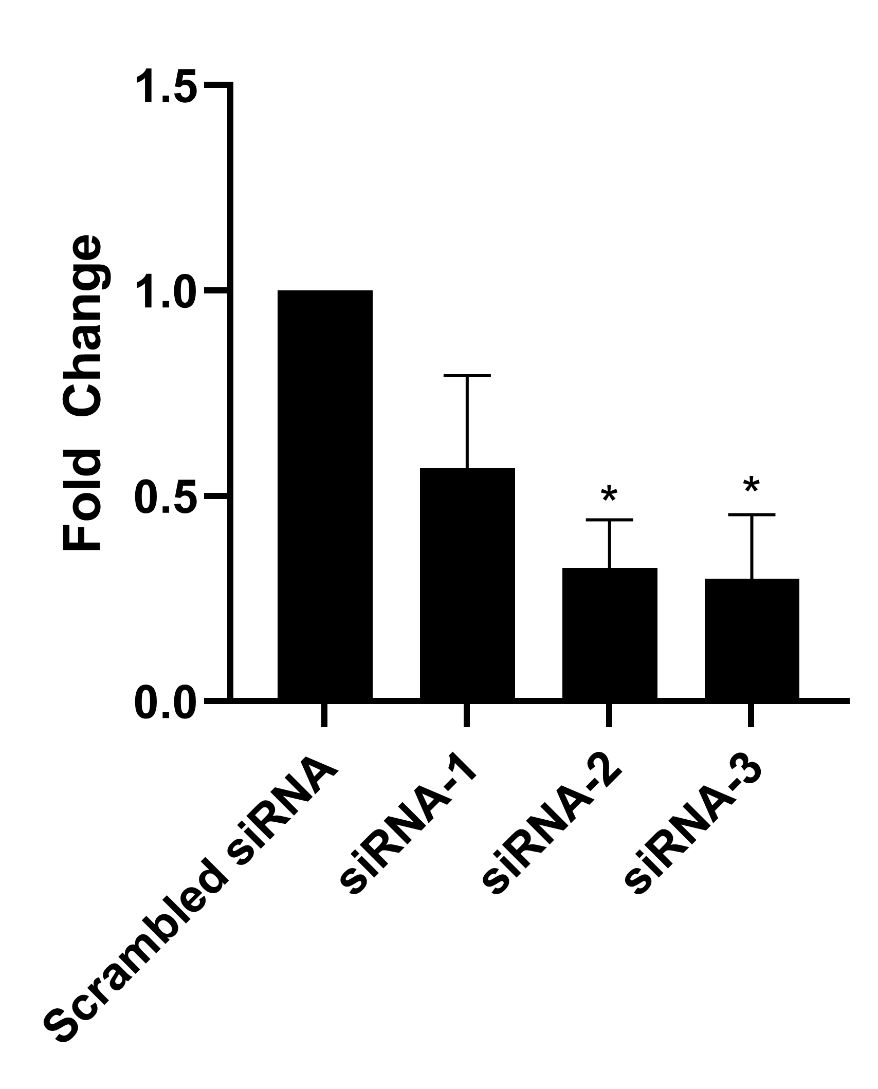

**Supplemental Materials 6. Tim-3 knock down by siRNA treatment**

Three *Tim-3* specific siRNA were tested for the ability to knock down expression in THP-1 differentiated macrophages as compared to cells treated with scrambled siRNA as negative control. siRNA-3 knocked down *Tim-3* expression by 70% (p = 0.0462, N = 3) and was the siRNA used in subsequent experiments. N = 3 (minimum). Mean ± SEM, *P<0.05.
