## Supplementary figures and images for "Secreted filarial nematode galectins modulate host immune cells"

### Supplemental Materials 3

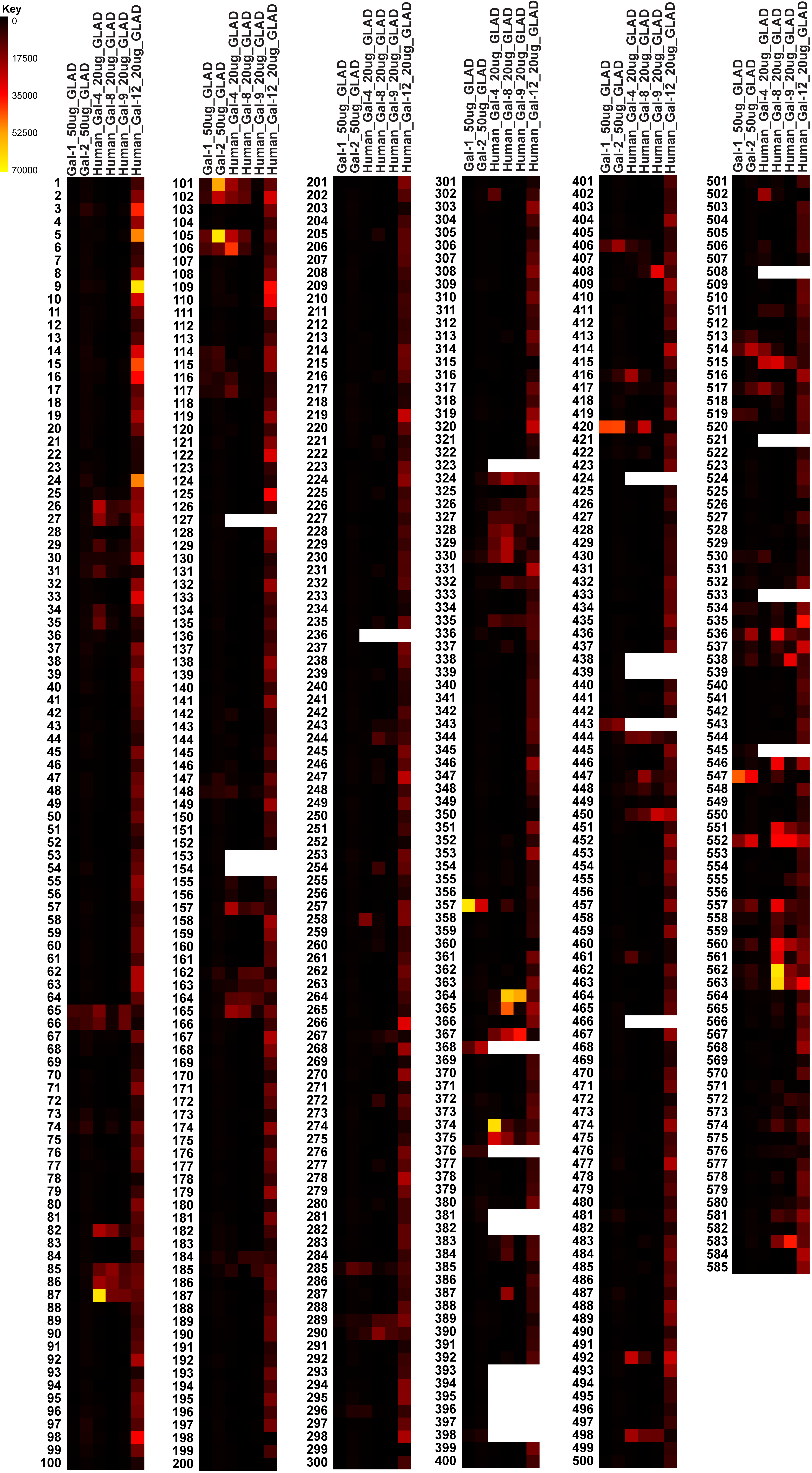
